# A real-time biochemical assay for quantitative analyses of APOBEC-catalyzed DNA deamination

**DOI:** 10.1101/2024.05.11.593688

**Authors:** Christopher A. Belica, Michael A. Carpenter, Yanjun Chen, William L. Brown, Nicholas H. Moeller, Ian T. Boylan, Reuben S. Harris, Hideki Aihara

## Abstract

Over the past decade, the connection between APOBEC3 cytosine deaminases and cancer mutagenesis has become increasingly apparent. This growing awareness has created a need for biochemical tools that can be used to identify and characterize potential inhibitors of this enzyme family. In response to this challenge, we have developed a Real-time APOBEC3-mediated DNA Deamination (RADD) assay. This assay offers a single-step set-up and real-time fluorescent read-out, and it is capable of providing insights into enzyme kinetics and also offering a high-sensitivity and easily scalable method for identifying APOBEC3 inhibitors. This assay serves as a crucial addition to the existing APOBEC3 biochemical and cellular toolkit and possesses the versatility to be readily adapted into a high-throughput format for inhibitor discovery.

## Introduction

The human APOBEC3 (A3) family is composed of 7 enzymes (A3A, B, C, D, F, G, and H) involved in the restriction of exogenous nucleic acids. The A3 proteins play important roles in innate immunity against viruses and transposons by catalyzing hydrolytic deamination of cytosine into uracil (C-to-U) in single-stranded (ss)DNA to inactivate the foreign genetic elements by hypermutation. However, recent evidence shows that at least two A3 members (A and B) are capable of targeting cellular genomic DNA in human cancers to elicit C-to-T and C-to-G mutations in preferred trinucleotide contexts (TCA and TCT), resulting in “APOBEC signature” mutations prevalent in over half of tumor types. Over the past decade, the expression of A3A/B and/or the presence APOBEC signature mutations has been associated with tumor development, metastasis, drug resistance, and poor clinical outcomes (1–4). These associative studies have been supported by work in mice where expression of catalytically active human A3A or A3B triggers APOBEC signature mutations and elevated tumor loads (5–8).

These clinical and animal experiments combine to suggest that inhibiting the A3A and A3B activities has the potential to improve the effectiveness of conventional anti-cancer therapies by limiting tumor evolution and preventing adverse outcomes such as drug resistance and metastasis. However, despite the significant advances made toward understanding the mechanisms by which A3A and A3B drive tumorigenesis and cancer progression, less progress has been made toward therapeutics designed to directly target and inhibit the mutagenic activity of these enzymes. One of the key factors preventing the rapid identification of small molecule inhibitors of A3A/B has been the lack of real-time assays to rapidly assess the enzymatic activity.

A commonly used method of analyzing A3 deaminase activities *in vitro* is a gel-based oligonucleotide cleavage assay (9–11). This assay has traditionally taken the form of a two-stage enzymatic reaction, in which an A3 enzyme is combined with a fluorescently labeled oligonucleotide containing the preferred sequence motif targeted by the A3 protein (*e*.*g*., 5’-TC motif for A3A and A3B, 5’-CC for A3G). After deamination of the target cytosine into uracil, the DNA is treated with a second enzyme, uracil-DNA glycosylase (UDG), to convert the 2’-deoxyuridine into an abasic site. The resulting DNA is treated with heat and/or a strong base, which causes a cleavage of the DNA backbone. As an alternative to the UDG treatment and heat/alkaline-mediated cleavage, we recently showed that *Pyrococcus furiosus* endonuclease Q (EndoQ) can be used to directly cleave DNA adjacent to 2’-deoxyuridine generated by A3B (12, 13). The readout of this assay is fluorescence-based quantification of both the cleaved and uncleaved DNA, separated by gel electrophoresis. Though reliable, this procedure does not provide a real-time readout to monitor the reaction kinetics and lacks the throughput required for large-scale screening of inhibitors. An NMR-based method allows real-time monitoring of the C-to-U conversion but requires much larger quantities of material and suffers from low throughput and high expense (9–11).

Here we report the development of a simple *in vitro* method for monitoring A3-mediated DNA deamination in real-time. This novel method only requires a single A3-containing reaction with a large excess of EndoQ and a self-quenched ssDNA reporter substrate, which allows sensitive detection of A3-mediated DNA deamination via a fluorescent plate reader or a gel-based read-out. The assay is both flexible and sensitive, making it a perfect candidate for usage in multi-well plates and high-throughput screens.

## Results

### The RADD assay detects A3B-mediated deamination in real-time

A drawback to most current methods of quantifying A3-mediated ssDNA deamination is that reactions are necessarily multi-step and usually involve the stepwise set-up of a reaction by first adding the deaminase to catalyze C-to-U, second adding UDG to remove the U, and finally adding NaOH/heat to break the backbone at the abasic site (the original site of deamination) (14) (schematic in **Fig. 1A**). Some studies have reduced this 3-step reaction to a 2-step version by adding a molar excess of UDG (15), but the requirement for a backbone cleavage step has still remained. An additional consideration for the implementation of such an assay on mechanized screening platforms is the potentially corrosive nature of NaOH stock solutions.

**Figure 1.**
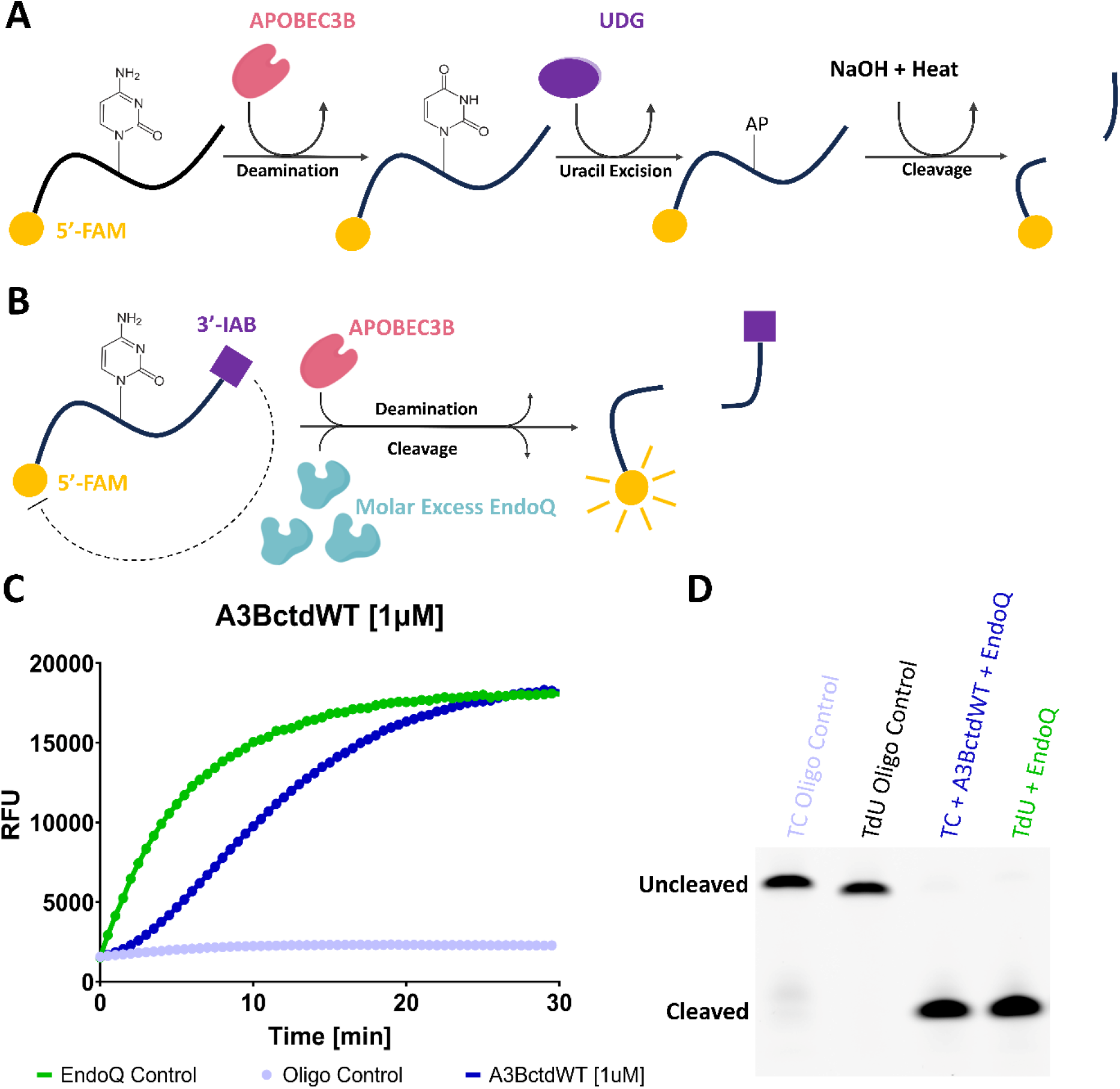
Fluorescence and gel-based readout of the RADD assay. **(A)** Schematic representing the molecular process involved in traditional UDG-based deamination assays. **(B)** Schematic of the molecular process which results in real-time fluorescence reporting of APOBEC deamination in a single step through the RADD assay. **(C)** Fluorescence readout of a RADD assay monitoring the deamination activity of 1 µM A3BctdWT. The A3BctdWT reaction (dark blue) contained 1 µM TC reporter and 2 µM EndoQ. The EndoQ control reaction (green) contained A3BctdWT protein storage buffer, 2 µM EndoQ, and 1 µM TdU reporter. The negative control (light blue) contained 2 µM EndoQ, A3BctdWT protein storage buffer, and 1 µM TC reporter. The EndoQ control highlights that EndoQ does not act as the rate-limited enzyme. **(D)** Gel readout of the reactions shown in C. Both the A3BctdWT and EndoQ control reactions show complete substrate processing within 30 minutes.

We sought to make this a single-step reaction and eliminate NaOH by combining the A3B-mediated deamination and endonuclease cleavage steps of the gel-based assay into a single step while utilizing a Förster resonance energy transfer (FRET) reporter oligo comprised of a 15nt 5’ 6-carboxyfluorescein (FAM) labeled ssDNA containing a 3’ Iowa Black® FQ (IAB) quencher (schematic in **Fig. 1B**). At the center of this reporter is a TCA motif which, when the cytosine is deaminated, is rapidly cleaved by a molar excess of EndoQ. Upon cleavage, the 5’-FAM is freed from its quencher allowing for a significantly increased fluorescence signal (**Fig. 1C**). This reporter was chosen based on our previous work which showed the sequence was readily deaminated by A3B and cleaved by EndoQ (12). Furthermore, the length of the reporter fell below the generally recommended maximum of 20-24nt for self-quenched fluorescent DNA probes (16, 17). To ensure that A3B is the rate-limiting enzyme, we have determined the kinetic parameters of EndoQ to show that the concentration of EndoQ used (2 µM) is >30-fold higher than its apparent dissociation constant (K_m_) for a dU-containing DNA probe (64 nM; Fig. S1A, B). We also confirmed that 2 µM EndoQ is saturating for 100 nM A3BctdWT by comparing the initial velocity of the fluorescence signal increase over a series of EndoQ concentrations (Fig. S1C, D). Additionally, since this assay was built off the foundation of the gel-based assay, the reactions can be run on a gel after completion to ensure rigor in the real-time data (**Fig. 1D**).

We next compared three different enzymatically active A3B constructs. The first was the A3B C-terminal catalytic domain containing a double-mutation for improved solubility L230K/F308K (A3BctdDM), whereas the second was the same C-terminal domain without mutations (A3BctdWT). The third construct was the full-length A3B purified from human cells (flA3BWT). Each of these enzymes was compared to a baseline control reaction containing an enzymatically inactive E255A derivative of the respective construct. In parallel, an EndoQ reaction containing the reporter oligo with a TdUA motif allowed for cleavage without A3B to confirm that EndoQ is not rate-limiting under these conditions. From these reactions, we were able to readily discern that each construct had different rates of deamination. Surprisingly, despite being the most soluble of all three enzymes, A3BctdDM (**Fig. 2A**) showed less activity than A3BctdWT (**Fig. 2B**). Both these enzymes showed significantly lower activity compared to flA3BWT (**Fig. 2C**). The results observed in the real-time fluorescence readings were also well supported by subsequent gel scans, showing a partial cleavage of the TC oligo by A3BctdDM and A3BctdWT, and slightly improved cleavage by flA3BWT (**Fig. 2D, 2E, 2F**).

**Figure 2.**
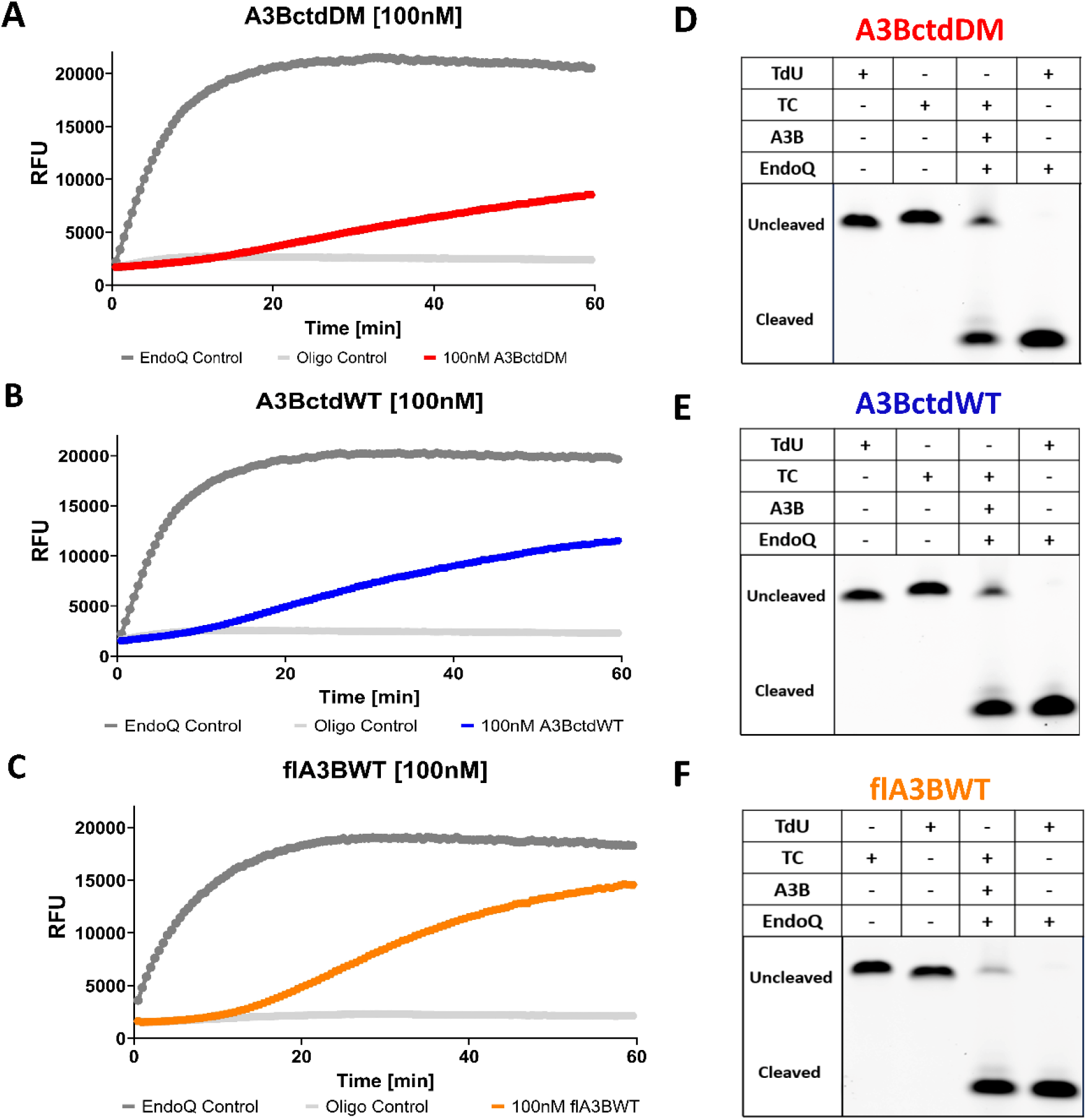
Comparative RADD assay readout for A3BctdDM, A3BctdWT, and flA3BWT. **(A-C)** The fluorescent signal of 100 nM A3BctdDM (Red), A3BctdWT (Blue) and flA3BWT (Orange) over the course of a 1-hour reaction containing 1 µM reporter oligo is shown. Each reaction is compared to an EndoQ control reaction containing 2 µM EndoQ and 1 µM TdU substrate. The negative control for each reaction contained 1uM TC reporter and 2 µM EndoQ, and A3B was substituted with the respective protein storage buffer. **(D-F)** Gel visualization of reactions from each assay. After fluorescence scanning, the assay samples were run on a TBE-urea gel to observe the relative substrate processing of each enzyme. The full-length 15nt oligo is the top band and the deaminated and cleaved substrate appears as the lower band. These results highlight the sensitivity of the fluorescence scan compared to the gel-only method.

### APOBEC3B enzyme kinetics determined using the RADD assay

Based on these initial results, we began testing these enzymes against a dilution series of the TC reporter in order to determine kinetic parameters. Similar to the initial experiments, 100 nM of either A3BctdDM, A3BctdWT, or flA3BWT was reacted with a series of increasing concentrations of substrate (Fig. S2). Consistent with initial observations, A3BctdDM was the slowest at processing the substrate with a turnover rate (K_cat_ = 0.0051 s^-1^) (**Fig. 3A**) similar to a previously reported value (18, 19). Interestingly, the K_m_ value for A3BctdDM determined from these experiments (4.4 µM) was substantially lower than the previously reported value of 320 µM (18, 19). A3BctdWT showed a slow, yet improved turnover rate compared to A3BctdDM as predicted (K_cat_ = 0.0059 s^-1^), and the K_m_ was determined to be 2.9 µM (**Fig. 3B**). Following a similar pattern, flA3BWT showed an improved turnover rate compared to A3BctdDM and A3BctdWT (K_cat_ = 0.0065 s^-1^), however, the observed apparent substrate affinity was nearly identical to that of A3BctdWT with the K_m_ being 3.0 µM (**Fig. 3C**). The data for all three constructs showed an excellent fit to the *Michaelis-Menten* model (**Fig. 3D**). Taken together, these results demonstrated that this assay can be used to reliably characterize the steady-state enzyme kinetics of different A3B constructs, and could potentially be used to assess the effects of different buffers or inhibition by small molecules.

**Figure 3.**
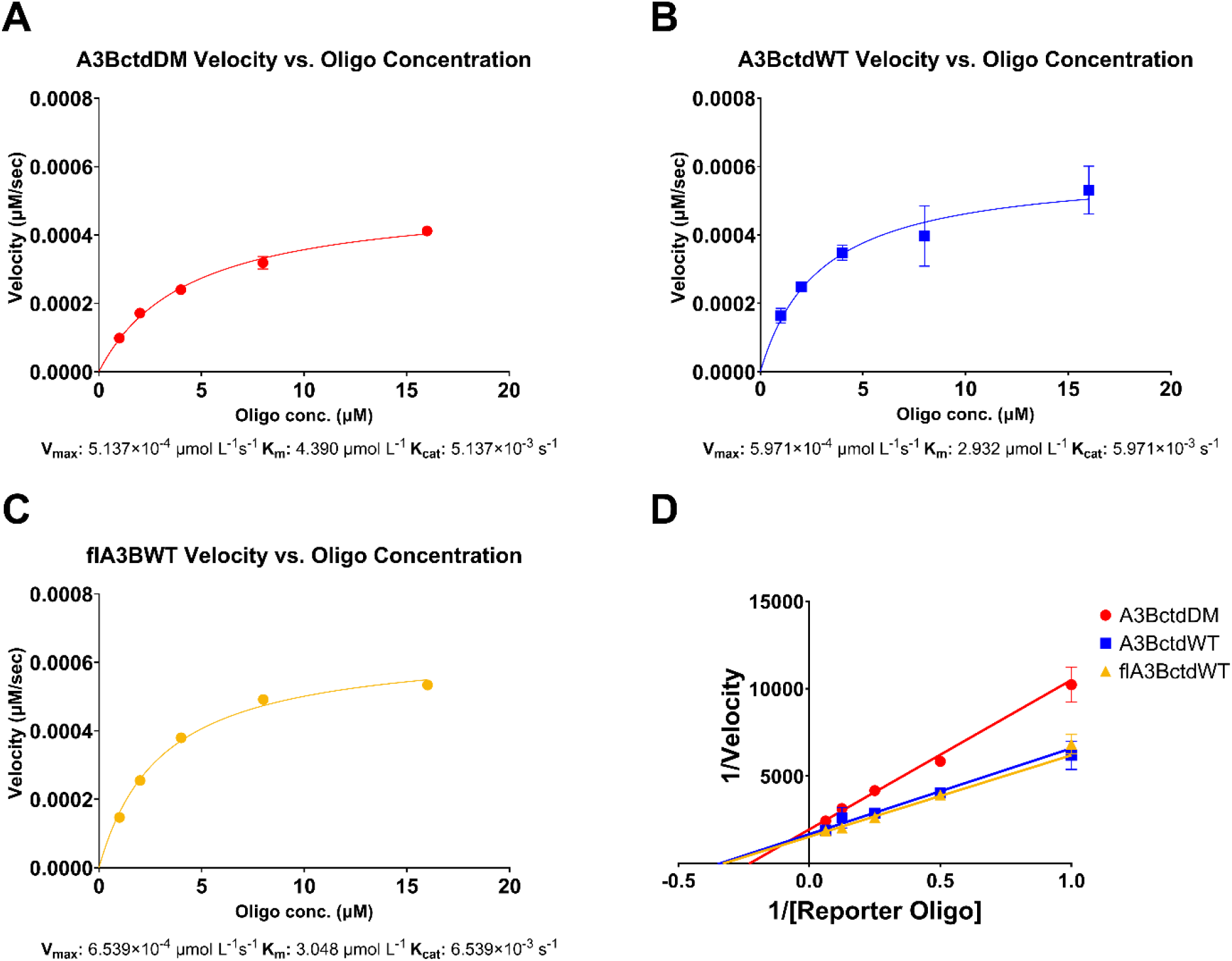
Enzyme kinetic parameters measured with the RADD assay. **(A)** Enzyme velocity of 100 nM A3BctdDM versus a ½ dilution series of 16 µM down to 1 µM reporter oligo (N = 3) fitted to the *Michaelis-Menten* model (R^2^ = 0.99). The predicted V_max_ was 5.1397×10^−4^ μmol L^-1^s^-1^ 95% CI [4.789×10^−4^ to 5.532×10^-4^], resulting in a K_cat_ of 5.137×10^−3^ s^-1^. The K_m_ of A3BctdDM was predicted to be 4.390 μmol L^-1^ 95% CI [3.651 to 5.290]. **(B)** Enzyme velocity of A3BctdWT versus the same reporter oligo dilution series described above (N = 3), fitted to the *Michaelis-Menten* model (R^2^ = 0.87). The predicted V_max_ for A3BctdWT was 5.971×10^-4^ μmol L^-1^s^-1^ 95% CI [5.051×10^−4^, 7.204×10^−4^], resulting in a K_cat_ of 5.971×10^−3^ s^-1^. The K_m_ was predicted to be 2.932 μmol L^-1^ 95% CI [1.723, 4.943]. **(C)** Enzyme velocity of flA3BWT versus the same reporter oligo dilution series described above (N = 3), fitted to the *Michaelis-Menten* model (R^2^ = 0.99). The predicted V_max_ for flA3BWT was 6.539×10^−4^ μmol L^-1^s^-1^ 95% CI [6.229×10^−4^, 6.873×10^−4^], resulting in a K_cat_ of 6.539×10^-3^ s^-1^. The K_m_ was predicted to be 3.048 μmol L^-1^ 95% CI [2.644 to 3.510]. **(D)** *Lineweaver-Burk* plot representing the enzyme kinetics of A3BctdDM, A3BctdWT, and flA3BWT.

An alternative method for estimating the deamination rate of A3B is to run the RADD assay in a single-turnover condition, using an A3B concentration well above the K_m_ and in excess over the reporter substrate. Such conditions ensure that the majority of the substrate is bound to the enzyme at any given time, allowing k_obs_ to be approximately equivalent to k_cat_. To test this methodology, a reaction using the standard 2 µM of EndoQ, an excess of 5 µM A3BctdDM, and 1 µM TC reporter was performed (Fig. S3). Under these enzyme concentration conditions, the first-order rate constant for the deamination reaction can be obtained using a mathematical model of sequential (consecutive) biochemical reactions (20, 21). First, the apparent rate constant for EndoQ (0.00297 s^-1^) was determined from the TdU controls in this experiment (**Fig. 4A**). This value was then used as k_2_ (the rate constant for the second step) in the sequential enzymatic reaction equation to fit the A3BctdDM data and estimate k_1_ (the rate constant for the first step) to be 0.00550 s^-1^ **(Fig. 4B, 4C)**, which aligns well with the k_cat_ determined using the *Michaelis-Menten* model and falls within the 95% confidence interval of that measurment. To verify the utility of this method, the sequential reaction model was also applied to intedpendant experimental data obtained using 1 µM A3BctdDM (**Fig. 4D**). Since the concentration of A3BctdDM is below K_m_, the rate measured does not directly translate to k_cat_. Regardless, the data show an excellent fit to the model and yielded an apparent first-order rate constant k_1_ of 0.00099 s^-1^.

**Figure 4.**
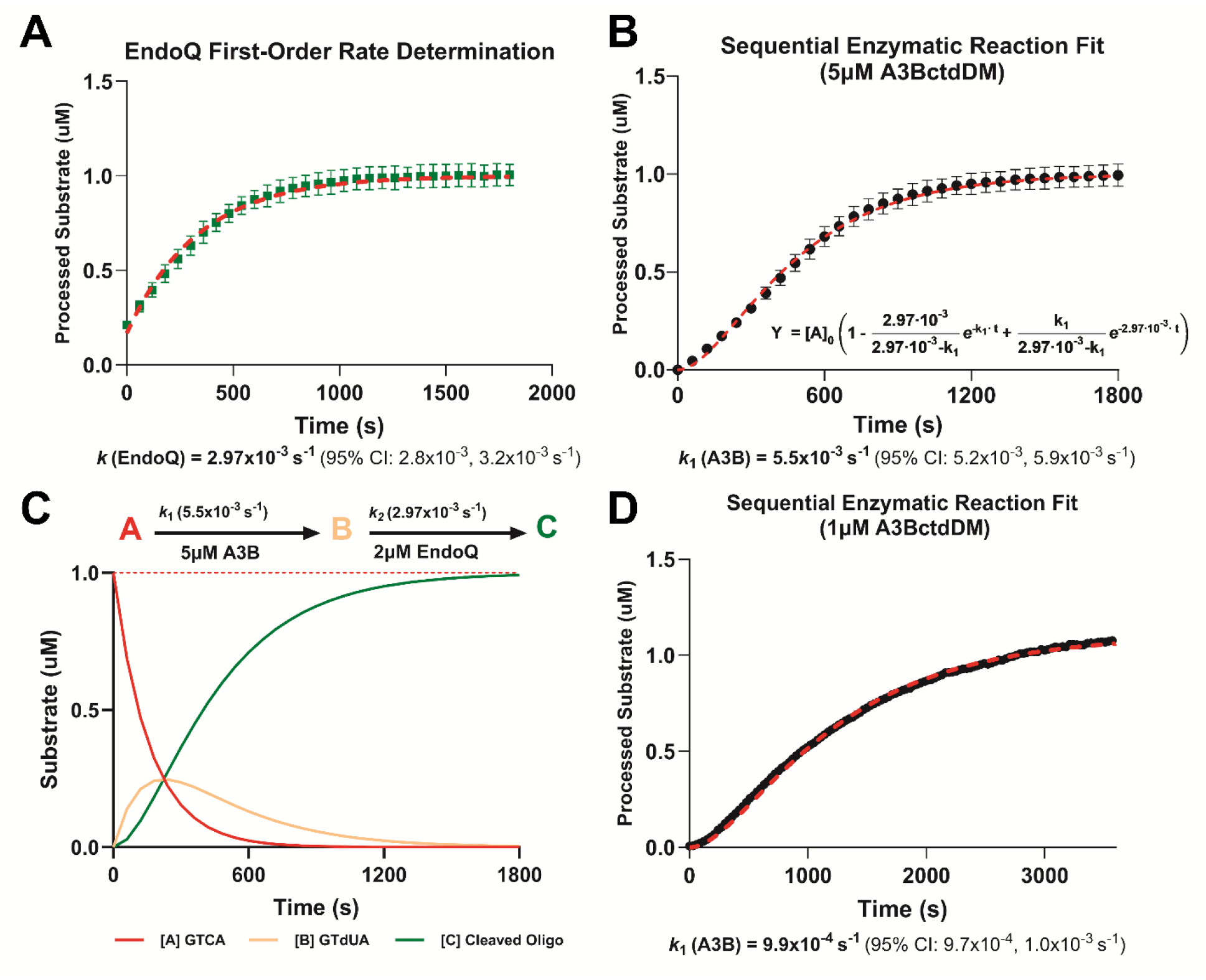
Estimating First-Order Rate Constants Using RADD and Sequential Enzymatic Reaction Modeling. **(A)** Estimating the k_cat_ of EndoQ by measuring the k_obs_ of EndoQ in a single-turnover condition (N = 3). The k_cat_ of EndoQ was determined to be 2.97x10^−3^ s^-1^ 95% CI [2.8x10^−3^ to 3.2x10^−3^] (R^2^ = 0.97). **(B)** Estimating the k_cat_ of A3BctdDM by measuring the deamination k_obs_ at 5 μM A3BctdDM (N = 3). Using the k_cat_ of EndoQ as k_2_ in the sequential enzyme reaction equation, the model showed an excellent fit to the A3BctdDM data (R^2^ = 0.98) and predicted a k_cat_ of 5.5x10^−3^ s^-1^ 95% CI [5.2x10^−3^ to 5.9x10^−3^ s^-1^]. **(C)** Schematic of theoretical concentrations of the TC reporter, deaminated reporter (TdU), and cleaved reporter during the 5 μM A3BctdDM deamination reaction simulated based on the rate constants determined in this experiment. **(D)** Estimating the k_obs_ of A3BctdDM by measuring the deamination rate of 1 μM A3BctdDM (N = 1). Using the same rate constant for EndoQ as above, the 1 μM A3BctdDM data was fitted with the sequential enzyme reaction equation (R^2^ = 0.99) to predict a k_obs_ of 9.9x10^−4^ s^-1^ 95% CI [9.7x10 to 1.0x10^−4^ s^-1^].

### The RADD assay Is a valuable tool for discovering APOBEC3B inhibitors

An important parameter when evaluating the efficacy of a real-time enzymatic assay for potential drug discovery is the Z’-factor, which indicates the likelihood of false negatives and positives by way of measuring the separation between positive and negative signals. A Z’-factor of ≥0.5 is considered an excellent assay metric for potential high-throughput screening (22). Using 100 nM A3BctdWT and an end point fluorescence reading at 1 hour, we were able to achieve a Z’-factor of 0.73, a signal-to-noise ratio of 76, and a signal-to-background ratio of 4.8 (**Fig. 5A**). All these factors combined to show that the assay provides a strong separation between positive and negative signals and support its potential value in high-throughput screening for anti-A3B compounds.

**Figure 5.**
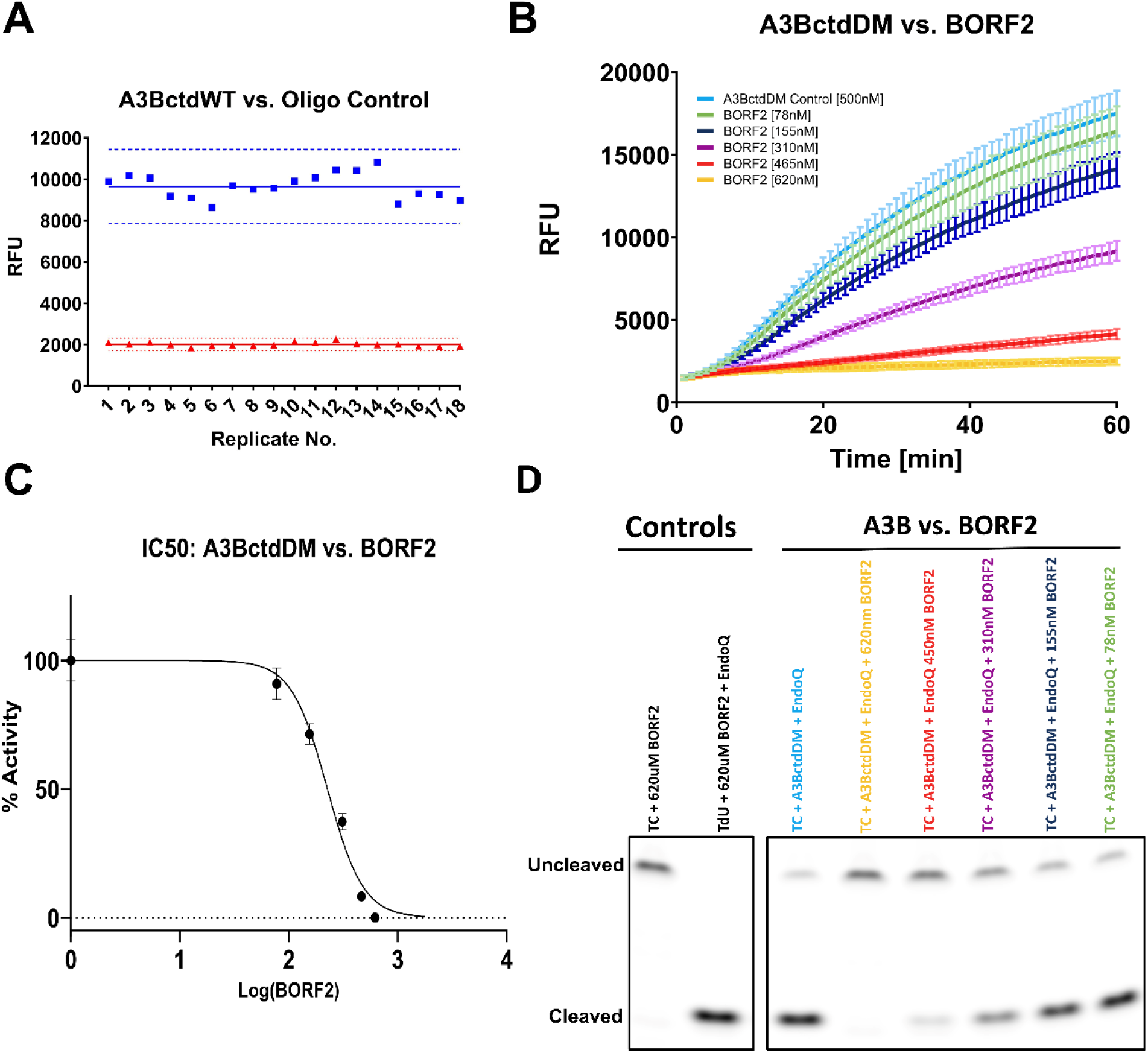
Measuring deaminase inhibition using the RADD assay. **(A)** The variation between positive (blue) and negative (red) fluorescence signals of a standard 1-hour reaction using 100 nM A3BctdWT (N = 18). The solid lines represent the mean endpoint signal while the dashed lines represent the mean ± 3×SD. The Z-factor of this assay was determined to be 0.73 while the signal-to-noise ratio was found to be 76, and the signal-to-background ratio was found to be 4.8. **(B)** The fluorescence readout of 500 nM A3BctdDM versus a dilution series of EBV BORF2 between 620 nM and 78 nM (N = 3). **(C)** IC50 graph displaying the percent-activity of 500nM A3BctdDM versus the concentration of BORF2 (R^2^ = 0.98). The top and bottom of the curve were constrained to 100 and 0 %, respectively. The IC50 of BORF2 was predicted to be 226.4 nM, 95% CI [205.1 to 249.1]. **(D)** The gel readout of the BORF2 inhibition assay confirms decreased substrate processing proportional to BORF2 concentration.

We further validated the assay using a potent natural inhibitor of A3B, the Epstein-Barr virus (EBV) large ribonucleotide reductase subunit BORF2. This viral protein is known to strongly bind and inhibit the catalytic activity of A3B as a means to protect the viral genome from A3B-mediated hypermutation (23, 24). The activity of A3BctdDM was measured against a dilution series of BORF2 to determine how effective our assay was at detecting inhibition. This set-up utilized 500 nM A3BctdDM for improved signal, and was incubated in the presence of 78, 155, 310, 450, and 620 nM BORF2. Based on the previous characterization of the A3B-BORF2 interaction with a low-nM K_D_, it was predicted that complete inhibition of A3B would be achieved by a stoichiometric concentration of BORF2, between 450 and 620 nM (24). For this assay, A3BctdDM and BORF2 were combined and incubated prior to addition to the remaining reaction components. A control reaction containing the TdU substrate, EndoQ, and 620 nM BORF2 showed that BORF2 does not interfere with the endonuclease activity of EndoQ. As an additional control to ensure that deamination was not affected by protein crowding, a similar dilution series of BSA was tested against A3B, showing no inhibitory effect (Fig. S4). The assay results showed a dose-dependent inhibition of A3BctdDM by BORF2, with the 620 nM BORF2 condition showing a complete ablation of the enzymatic activity (**Fig. 5B**, Fig. S5). These data were then used to calculate an IC50 value for BORF2, showing an approximate IC50 of 226 nM (95% confidence interval: 205.1 to 249.1 nM), fully consistent with the expected tight binding of BORF2 to A3B (**Fig. 5C, 5D**).

### The RADD assay can be used to detect APOBEC3B activity in cell lysate

Another use for the RADD assay we explored is the potential to detect active full-length A3B in human cell lysate. We began by transfecting HEK293T cells, which do not endogenously express A3B at detectable levels (25), with an HA-tagged full-length A3B expression vector. A control culture was transfected with an identical vector lacking the A3B coding sequence. These cells were then diluted to a concentration of 70,000 cells / μL of lysis buffer and lysed via sonication. The clarified lysate was then serially diluted 1:2 and 1:4 in lysis buffer. Rather than the standard TC and TdU reporters, this experiment utilized a modified reporter containing an internal ZEN quencher in addition to the standard 5’-Iowa Black quencher due to their enhanced signal/background (Fig. S6) and observed ability to better resist non-specific degradation. Using 2 µM EndoQ and 1 µM reporter, the real-time readout showed a clear signal from the A3B-containing reactions which decreased in a step-wise manner according to the dilution factor (**Fig. 6A**, Fig. S7) while the reactions containing control lysate showed no deamination activity. Furthermore, the gel-based readout from this assay mirrored the results observed in real-time fluorescence scanning (**Fig. 6B**). Taken together, these results show that the RADD assay is highly versatile and can be used to detect A3B deamination activity in human cell lysate.

**Figure 6.**
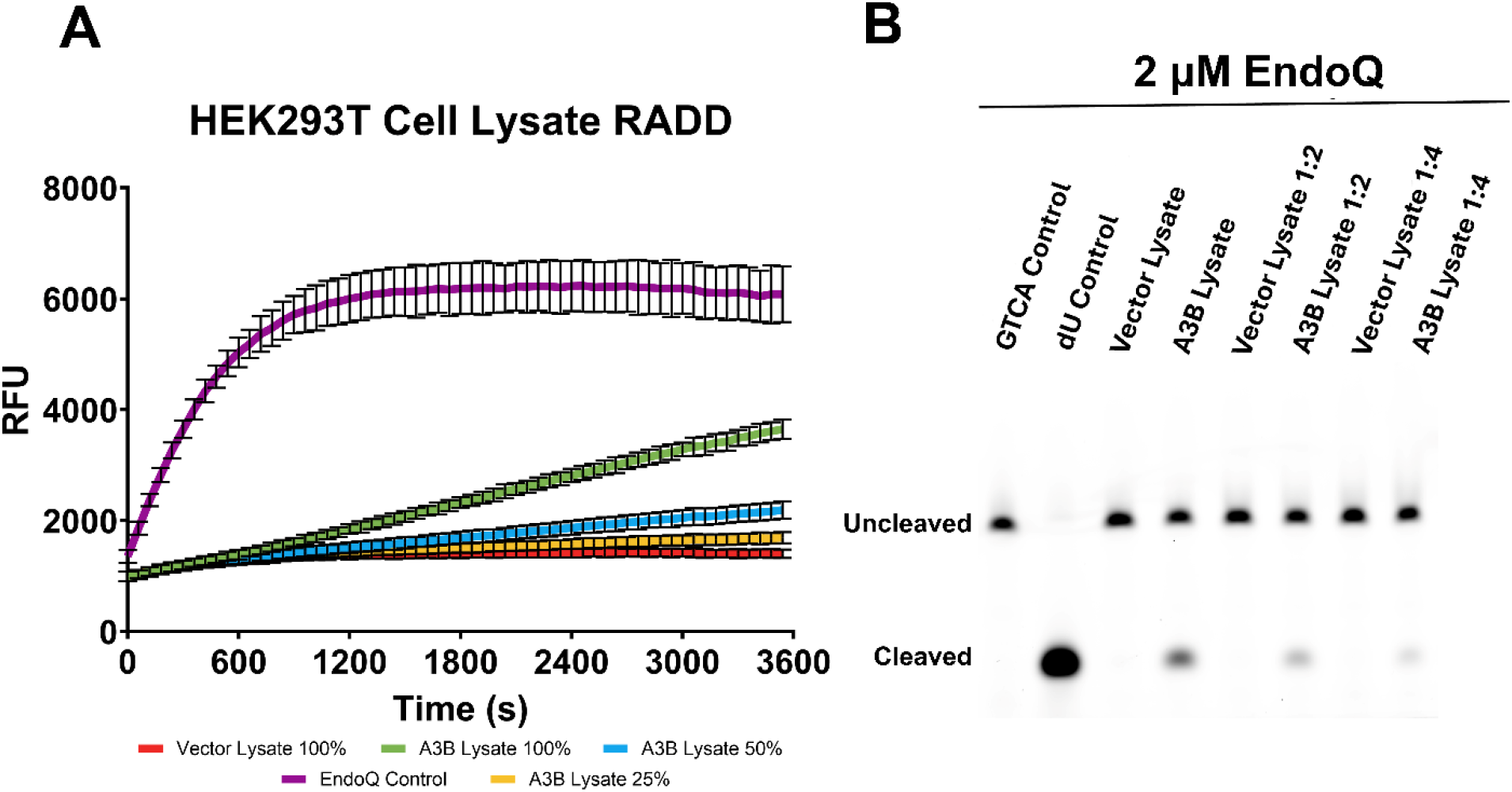
Measuring APOBEC3B activity in HEK293T cell lysate. **(A)** Real-time deamination readout of reactions containing various dilutions of A3B-expressing human cell lysate and controls transfected with empty vector (N = 3 for each condition). The trace in purple represents the EndoQ control reaction containing 1 μM TdU-ZEN, 2 μM EndoQ, and lysis buffer. The green, blue, and yellow traces represent 1 μM TC-ZEN reporter in the presence of undiluted, 1:2, or 1:4-diluted A3B-expressing cell lysate, respectively. The red trace represents the negative control, containing undiluted control cell lysate. All reactions contained 2 μM EndoQ. **(B)** After fluorescence scanning, the reactions were run on a 15% TBE-Urea gel. None of the control cell lanes showed any substrate processing (lower band) while the A3B-containing samples showed decreased substrate processing in-line with increased dilution.

## Discussion

Here we have shown that the RADD assay is a useful tool for measuring A3B enzyme kinetics and inhibition in real-time. This assay benefits from single-step simplicity while still providing sensitive real-time data much more quickly than biochemical or cell-based counterparts. We outline two methods of determining APOBEC enzyme kinetics using the RADD assay. The first being the *Michaelis-Menten* model which allows for the determination of K_m_, V_max_, and k_cat_, and the second being the sequential/consecutive biochemical reactions model which can be used to determine k_obs_ or k_cat_ depending on the enzyme concentration in relation to the K_m_ for a given substrate. While the *Michaelis-Menten* model can be used to identify a wider range of kinetic variables, the consecutive biochemical reactions model offers a more rapid determination of the first-order rate constant and may be a marginally more accurate estimation as it takes the consecutive nature of the RADD reaction into account. One interesting observation made during assay development was the difference in enzymatic activity observed between A3BctdDM, A3BctdWT, and flA3BWT. The lysine mutations made to A3BctdDM (L230K/F308K) are located more than 15 Å away from the active site of the enzyme and are not directly involved in DNA binding or catalysis. Thus, the slightly lower deaminase activity of A3BctdDM, despite its better solubility and much lower propensity to aggregate in solution, could be due to subtle changes in the tertiary structure of the protein having an allosteric effect on the substrate binding or catalysis. The N-terminal domain (NTD) of A3B has a similar deaminase fold as A3Bctd but does not have enzymatic activity, and its role remains poorly understood. Our kinetic analyses showed that flA3BWT has a similar substrate affinity, but improved catalysis (similar K_m_ despite higher K_cat_). These observations suggest that the presence of the NTD may have additional allosteric effects that enable slightly improved catalysis. Furthermore, it appears that the NTD has little effect on substrate binding, however, this may be a result of the short length of or the location of the TCA within the reporter oligo, as previous research has observed improved substrate cleavage of long substrates compared to short substrates using full-length A3B (26).

The kinetic parameters we obtained for A3BctdDM measured via the *Michaelis-Menten* model and the consecutive biochemical reactions model generally agree with previously reported values, with a greater variation in K_m_ (18). This is likely caused by different buffer conditions such as ionic strength, which we observed can significantly affect deamination activity. Differences in the length/sequence of ssDNA substrates may also be influential. For instance, the presence of G at the –2 position in our reporter may have positively affected deamination by A3B (26). It would be interesting to systematically examine the substrate preferences of A3B using the RADD assay. The K_m_ of 4.4 µM observed in our experiments is a realistic estimate of A3BctdDM affinity to our substrate based on the previously observed affinity of A3Bctd for ssDNA of an identical length (27). This assay is fully compatible with other TC-preferring A3 enzymes, including A3A, and with A3 members with different sequence preferences by simply changing the DNA sequence of the substrate. Furthermore, given the ability of EndoQ to efficiently cleave DNA strands with either dU or deoxyinosine, the assay can readily be adapted to characterize the activities of DNA adenine deaminases (28).

The RADD assay is likely to become a useful addition to the A3 biochemical toolbox (29), in part because it can be easily translated to a high-throughput format due to the plate-based nature of the assay and its high Z’-factor. Additionally, we have found that the assay is tolerant of at least 10% DMSO without any negative impact on the rate of deamination (Fig. S8), making the prospect of using this assay for small molecule inhibitor testing even more straightforward. By running each experiment with a corresponding TC, TdU, and TdU + EndoQ control, nuclease contamination, compounds that inhibit EndoQ, and naturally fluorescent compounds can easily be detected and thereby minimize the potential for false positives and effects of other confounding factors. The BORF2 inhibition assays performed here further suggest that this novel assay is capable of sensitively detecting A3B inhibition and accurately determining IC50 values.

In addition to using purified deaminases, we showed that RADD can be used to detect A3B activity in human cell lysate, which allows for examination of DNA deaminase activities in more native-like conditions without the need for purification. It should be noted that the deamination activity detected via this method is generally lower compared to that of purified proteins. This observation could be due to a combination of reduced protein concentrations and the presence of EDTA in the lysis buffer. EDTA is necessary to prevent non-specific nuclease activities, however, it also negatively impacts the activity of EndoQ and A3B which use Mg^2+^ and Zn^2+^, respectively, as essential co-factors. While the activity could be decreased, we showed that the signals produced by reporter cleavage are APOBEC-specific. The RADD assay with cell lysate provides a valuable platform for semi-quantitative comparison between mutant enzymes and evaluating inhibitors.

The RADD assay can also be used in tandem with the recently developed assay for observing A3 deaminase activities in living cells, the APOBEC-mediated base editing reporter (AMBER) (30). This reporter plasmid contains a cytosine base editor (CBE) with corresponding guide RNA. The CBE is composed of either A3A or A3B fused to a Cas9 nickase and a uracil-DNA glycosylase inhibitor. Alongside the CBE, the plasmid contains a constitutively expressed mCherry for transfection normalization and eGFP with a missense (L202S) mutation that prevents fluorescence. When expressed, the guide RNA will recruit the CBE to the eGFP sequence and mutate the TCA (serine) codon to TTA (leucine), restoring eGFP fluorescence. This cell-based assay offers several advantages by allowing for higher throughput, real-time monitoring of APOBEC activity via fluorescence signal. The AMBER system also prevents false positives due to cytotoxicity and assures that positive hits are cell-permeable. On the other hand, this assay also rules out potentially effective A3B inhibitors that could be modified to gain cell permeability. Furthermore, the AMBER system provides a comparatively slow read-out, taking 48-72 hours to reach peak fluorescence. Though a powerful cell-based assay, the AMBER system lacks the flexibility of an *in vitro* assay and does not diminish the need to develop a complementary real-time biochemical assay for A3 activities such as the RADD assay described here.

Looking ahead, potential improvements to the RADD assay can be made by reducing the background fluorescence of the reporter substrate. It was observed that during the first 5 minutes of fluorescence scanning, there is a small increase in signal from the oligo control before reaching a stable baseline, despite no clear oligo degradation occurring. This effect is reduced by substrate pre-incubation but could not be eliminated completely. A potential method to remove this effect and reduce overall background fluorescence is the usage of double-quenched reporters. A doubly quenched reporter containing an additional internal ZEN quencher has yielded a lower background and a better signal-to-noise ratio without negatively affecting the rate of deamination by A3B (**Fig. 6**, Fig. S6).

## Experimental procedures

### Standard fluorescent reporter oligos

The primary reporter oligo utilized in this assay (TC) is a 15 nucleotide (nt) ssDNA with a single target cytosine in a TCA motif, a 5’ 6-carboxyfluorescein (FAM), and a 3’ Iowa Black® FQ (IAB) quencher (5’-FAM-TAGGT<u>C</u>ATTATTGTG-IAB-3’). Position 6 of this ssDNA oligo is the only cytosine base and therefore the only site for A3-catalyzed deamination. The EndoQ control reporter (TdU) is nearly identical to the primary reporter with the exception of a 2’-deoxyuridine at position 6 rather than a 2’-deoxycytidine. All oligonucleotides were synthesized by Integrated DNA Technologies, Inc.

### Double-quenched fluorescent reporter oligos

The reporter oligos used for detecting A3B activity in human cell lysate have identical sequences to the singly quenched TC and TdU counterparts with the exception of an internal ZEN™ quencher between nt 9 and 10 (5’-FAM-TAGGT<u>C</u>ATT-**ZEN**-ATTGTG-IAB-3’). All oligonucleotides were synthesized by Integrated DNA Technologies, Inc.

### Protein purification

#### A3BctdDM / A3BctdWT

pET42b(+)GST-6xHis-A3BctdDM and pET42b(+)GST-6xHis-A3BctdWT plasmids, which contain codon-optimized genes for human A3B residues 187 to 378 with or without the solubility-enhancing mutations (L230K/F308K) fused to an N-terminal GST-6xHis tag with a HRV-3C protease cleavage site, were used to transform chemically competent C41(DE3) *E. coli*. The transformed cells were grown on an LB-agarose plate in the presence of 40 µg/mL kanamycin overnight at 37°C. All colonies growing on the plate were then resuspended and distributed to four 1-liter flasks of LB media containing 40 µg/mL kanamycin. These cultures were grown at 37°C until an OD of 0.8-1.0 was reached. Cultures were then induced with 0.5 mM isopropyl β-D-1-thiogalactopyranoside and 50 μM ZnCl_2_ at 18°C for 18 hours. The cultures were then centrifuged at 4,000xg for 30 minutes at 4°C. The pelleted cells were resuspended in 66 mL of buffer A (20 mM Tris-HCl pH 7.4, 500 mM NaCl, 5 mM imidazole, 5 mM β-mercaptoethanol).

Cells were lysed with the addition of 40 mg hen egg white lysozyme followed by sonication. The cell lysate was centrifuged at 64,000xg, at 4°C for 1 hour, followed by vacuum filtration through a 0.22 μm filter. The filtered lysate was then passed over a 5 mL Ni-NTA Superflow column (Qiagen). The column was washed extensively with buffer A and subsequently, the bound protein was eluted with a linear gradient of buffer A to B (20 mM Tris-HCl pH 7.4, 500 mM NaCl, 400 mM imidazole, 5 mM β-mercaptoethanol). Fractions containing GST-A3Bctd were pooled and treated with the HRV-3C protease to remove the GST-6xHis tag. The cleaved protein was then concentrated and underwent size exclusion chromatography using a HiLoad Superdex 75 26/600 pg column using A3B storage buffer (20 mM Tris-HCl pH 7.4, 200 mM NaCl, 1 mM tris(2-carboxyethyl)phosphine (TCEP)). Protein was snap-frozen in liquid nitrogen and stored at - 80°C.

#### MBP-BORF2(1-739)

Chemically competent BL21(DE3) *E. coli* was transformed with a pMalX(E)-BORF2(1-739) plasmid and grown on an LB-agarose plate containing 100 µg/mL ampicillin overnight at 37°C. Colonies were then resuspended and distributed to 4, 1-liter flasks of TB media containing 100 µg/mL ampicillin. These cultures were grown at 37°C until an OD of 0.8-1.0 was reached. Cultures were then induced with 0.5 mM IPTG at 18°C for 18 hours. The cultures were then centrifuged at 4,000xg for 30 minutes at 4°C. The cells were resuspended in 66 mL of a modified buffer A (20 mM Tris-HCl pH 7.4, 500 mM NaCl, 5 mM Imidazole, 5% glycerol, 5 mM β-mercaptoethanol). Cells were lysed with the addition of 40 mg lysozyme followed by sonication. The cell lysate was centrifuged at 64,000xg, at 4°C for 1 hour, followed by vacuum filtration through a 0.22 um filter. The filtered lysate was then passed over a 5 mL Ni-NTA Superflow column (Qiagen). Protein elution was carried out with a linear gradient of modified buffer B (20 mM Tris-HCl pH 7.4, 500 mM NaCl, 400 mM imidazole, 5% glycerol, 5 mM β-mercaptoethanol). The pooled protein was then concentrated and underwent size-exclusion chromatography using a Superdex 200 increase 10/300 column operating with BORF2 storage buffer (20 mM Tris-HCl pH 7.4, 200 mM NaCl, 5% glycerol, 1 mM TCEP). Protein was snap-frozen with liquid nitrogen and stored at -80°C.

#### Full-length-PfuEndoQ

A pET24-based expression plasmid for C-terminally 6xHis-tagged full-length *Pyrococcus furiosus* EndoQ (12) was used to transform chemically competent BL21(DE3) E. coli, which were grown on an LB-agarose plate in the presence of 40 µg/mL kanamycin overnight at 37°C. Obtained colonies were then resuspended and distributed to four 1-liter flasks of LB media containing 40 µg/mL kanamycin. These cultures were grown at 37°C until an OD of 0.8-1.0 was reached. Cultures were then induced with 0.5 mM IPTG and 50 μM ZnCl_2_ at 18°C for 18 hours. The cultures were then centrifuged at 4,000xg for 30 minutes at 4°C. The cells were resuspended in 66 mL of buffer A. Cells were lysed with the addition of 40 mg lysozyme followed by sonication. The cell lysate was centrifuged at 64,000g, at 4°C for 1 hour, followed by vacuum filtration through a 0.22 μm filter. The filtered lysate was then passed over a 5 mL Ni-NTA Superflow column (Qiagen). Protein elution was carried out with a gradient of buffer B. Pooled fractions containing PfuEndoQ were concentrated and underwent size-exclusion chromatography using a HiLoad Superdex 75 26/600 pg column operating with the A3B storage buffer. The purified protein was snap-frozen with liquid nitrogen and stored at -80°C.

#### Full-length A3B (WT)

A 200 mL culture of Expi293F cells (Thermo Fisher) was transfected, following the standard Expi293 transfection protocol, with pcDNA3.1-hsA3Bi-mycHis adding the transfection enhancers 18 hours post-transfection. Cells were harvested three days post-transfection by centrifugation for 5 minutes at 1000 g then the cell pellet was frozen at -80 °C. Cells were thawed on ice then resuspended in 50 mL lysis buffer (25 mM Tris-HCl pH 8.0, 5% glycerol, 0.1% IGEPAL CA-630, 20 mM imidazole, 2 mM magnesium chloride, and 0.5 mM TCEP). RNaseA was added to 100 µg/mL as well as 10 µL of Benzonase. Cells were sonicated on ice with a Branson Sonifier 450 for two 40% duty cycles of two minutes at power 5. The lysate was then incubated at 37 °C for an hour to allow the nucleases to work. Sodium chloride was added to a final concentration of 300 mM before centrifugation of the lysate at 16,000 g for 30 minutes at room temperature. The supernatant was collected, and sodium chloride was added to a final concentration of 1 M. NiNTA-sepharose (Qiagen) was added to the lysate followed by rotation for two hours at 4 °C. The chromatography media was collected by pouring the lysate through a Bio-Rad Poly-Prep column. The Ni-NTA sepharose was washed with 25 mM HEPES pH 7.4, 300 mM sodium chloride, 0.1% Triton X-100, 40 mM imidazole, and 10% glycerol. Elution was performed in 25 mM HEPES pH 7.4, 150 mM sodium chloride, 300 mM imidazole, 0.1% Triton X-100, 20% glycerol, and 1 mM TCEP. Protein was flash frozen in liquid nitrogen and stored at -80 °C.

### Enzyme stock preparation

A3BctdDM, A3BctdWT, and BORF2 were all prepared freshly from storage stocks for each experiment. Fresh storage stocks were thawed and centrifuged at 18,000xg at 4°C for 10 minutes to remove insoluble aggregates. Stocks were then quantified via UV absorbance measurement on a Nanodrop spectrophotometer blanked with the respective storage buffer. All enzymes were diluted into their respective storage buffer to a concentration 10X of the final concentration desired for the reaction. EndoQ reaction stocks were prepared by first thawing a fresh storage stock and heated at 60°C for 15 minutes to inactivate the nuclease activity of residual contaminants. The heat-treated stocks were then centrifuged at 18,000xg at 4°C for 10 minutes. The stock was then diluted to 20 μM in A3B storage buffer. These reaction stocks were then flash-frozen in liquid nitrogen and stored at -80°C.

### HEK293T cell culture and transfection

HEK293T cells (ATCC) (600,000 cells) were plated in a 100 mm cell culture dish. On the next day, they were transfected with 7.5 μg of A3B-HA_pcDNA3.1 (31), or pcDNA3.1_vector using TransIT-LT1 (MirusBio MIR2306). After 48 hours, the cells were washed with PBS/EDTA, released with Trypsin, and pelleted by centrifugation. The cell pellets were washed with PBS then frozen at -20°C.

### HEK293T cell lysis

Control and A3B-expressing HEK293T cells were thawed and diluted to ∼70,000 cells / μL in HED lysis buffer consisting of 25 mM HEPES pH 7.4, 15 mM EDTA, 10% glycerol, and Roche cOmplete™ EDTA-free protease inhibitor cocktail (Sigma-Aldrich). The diluted cells were then lysed in a sonicating water bath at 4°C for 20 minutes. After sonication the lysate was supplemented by PureLink™ RNase A (Thermo Fisher) to a concentration of 100 μg / mL and placed on a rotating shaker for 1 hour at room temperature. The lysate was then clarified by centrifugation at 18,000xg at 4°C for 10 minutes.

### Real-time deamination reactions

#### Standard deamination reactions

Each 40 μL reaction was carried out on a black 96-well half-area plate (Corning #3993). The 1X reaction buffer consisted of 60 mM Tris-HCl pH 8.0, 0.01% Tween 20, 1% DMSO, and 1 mM DTT. For general activity and kinetics assays, 24 μL of nuclease-free water was combined with 4 μL of 10X reaction buffer and 4 μL of 10 μM reporter. This mixture was pre-incubated at 37°C for 15 minutes while the enzyme mixture was prepared. The enzyme mixture consisted of 4 μL of 20 μM EndoQ and 4 μL of either A3B or A3B storage buffer. The enzyme mixture was also pre-incubated at 37°C for 10 minutes. After the pre-incubation step, the reaction mixture and enzyme mixture were combined in a well and fluorescence scanned for 1 hour at 37°C. For inhibition assays, the reaction mixture was comprised of 20 μL nuclease-free water, 4 μL 10X reaction buffer, and 4 μL of 10 μM reporter. The enzyme mixture consisted of 4 μL EndoQ, 4 μL A3B, and 4 μL BORF2 or BORF2 storage buffer. Pre-incubation and fluorescence scanning were performed identically to the standard activity assay.

#### Cell lysate deamination reactions

Similarly to the standard deamination reaction procedure, each 40 μL reaction was carried out on a black 96-well half-area plate. The 1X reaction buffer consisted of 60 mM Tris-HCl pH 8.0, 0.01% Tween 20, 1% DMSO, and 1 mM DTT. For cell lysate deamination assay, 18 μL of nuclease-free water was combined with 4 μL of 10X reaction buffer and 4 μL of 10 μM double-quenched reporter. This mixture was then combined with 10 μL clarified cell lysate and 4 μL 20 μM EndoQ. The reactions were then mixed by pipetting before transferring into the 96-well plate and scanned for 1 hour 37°C.

### Fluorescence scanning

All fluorescence readings were recorded with a TECAN Spark 10M instrument in concert with the SparkControl V3.0 software. Data were collected at 37°C using the scanning fluorescence kinetic mode, with the excitation wavelength and bandwidth set to 480 nm and 20 nm respectively. The emission wavelength was set to 520 nm and the bandwidth was set to 20 nm. Instrument gain was set to 71. The Z-position of the emitter was set to 16156 μm. All scanning sessions were recorded over 1 hour using either 30-second or 1-minute scan intervals.

### Gel-based read-outs

After real-time fluorescence scanning, reactions were combined in a 1-to-1 volume ratio with 100% formamide and heated to 95°C for 5 minutes to inactivate the enzymes. 12 μL of the inactivated reaction was loaded onto a 15% TBE-urea polyacrylamide gel (Invitrogen) and run at 300V for 35 minutes. Gels were scanned using an Amersham Typhoon imager and Amersham Typhoon Scanner Control Software 2.0.0.6 with the built-in Cy2 scanning method.

### Western blots

All samples were diluted 1:2 in reducing 2X SDS loading buffer and boiled at 95°C for 10 minutes. Samples were loaded onto a Mini-PROTEAN® TGX^™^ gel (Bio-Rad) and ran at 90V for 15 minutes followed by a subsequent run at 150V for 1 hour. Gels were then transferred at 80V for 1.5 hours at 4°C before blocking in PBST+4% milk overnight at 4°C. Blots were then incubated with primary rabbit α-HA antibody (Cell Signaling Technology #3724S) diluted 1:5,000 in PBST+4% for 2 hours at room temperature. This step was then followed by 6, 5-minute washes with PBST+4% milk. Blots were then incubated with fluorescently labeled secondary goat α-rabbit 680TL antibody (LI-COR Biosiences #925-68021), diluted 1:20,000 in PBST+ 4% milk + 0.01% SDS for 1 hour at room temperature. Blots were washed with PBST for 5 minutes a total of 6 times. Blots were imaged using an Amersham Typhoon imager and Amersham Typhoon Scanner Control Software 2.0.0.6 with the built-in IR-Short scanning method.

### Data analysis

#### A3B and EndoQ Michaelis-Menten kinetics

Kinetic analysis was performed by measuring the initial velocity of the enzyme reaction from the maximum slope of the fluorescence reading for each reporter concentration. Slope determination was performed using ICEKAT (32) after background fluorescence subtraction. The conversion of RFU-to-μM substrate was achieved by determining the maximum fluorescence of the fully processed 1 μM substrate and dividing the rate by this value to obtain the velocity in units of μM/sec. K_m_ and V_max_ were determined using the built-in *Michaelis-Menten non-linear fit* model in the GraphPad Prism software (Dotmatics).

#### Sequential enzymatic reaction model kinetics

EndoQ and A3B fluorescence intensities were both divided by the average maximum observed RFU value for their respective reporters to convert RFU to μM processed substrate. The k_obs_ for EndoQ was determined via the one-phase association model in Graphpad Prism, with the plateau constrained to 1.00. This value was then input into the user-defined equation seen above (Fig. 4B) and used to fit the data in GraphPad Prism with the “[A]_0_” variable (initial substrate concentration) constrained to 1.00.

#### BORF2 IC50

The BORF2 inhibition data were analyzed by determining the initial velocity via ICEKAT and translating the values into the percent-activity of the uninhibited reaction. The inhibitor concentration was translated to a log scale and IC50 was calculated via the built-in *log(inhibitor) vs. response – Variable slope* model in GraphPad Prism with constraints for the top and bottoms set to 100 and 0, respectively.

## Data availability

The data underlying this article are available in the Zenodo digital archive, at 10.5281/zenodo.11135133

### Supporting information

This article contains supporting information.

## Acknowledgments

We thank Dr. Daniel Harki and McKenzie Wyllie for their suggestions and discussion regarding statistical analysis and Ian Boylan for assistance during the setup of initial experiments.

## Funding and additional information

This work was supported by grants from the US National Institutes of Health (NIH), NCI P01CA234228 to R.S.H. and H.A and NIGMS R35GM118047 to H.A, and a Recruitment of Established Investigators Award from the Cancer Prevention and Research Institute of Texas CPRIT RR220053 to RSH. C.B. was supported by the UMN Institute of Molecular Virology and the US National Institutes of Health T32 training program NIH T32AI083196. R.S.H. is the Ewing Halsell President’s Council Distinguished Chair at University of Texas San Antonio and an Investigator of the Howard Hughes Medical Institute. The content is solely the responsibility of the authors and does not necessarily represent the official views of the National Institutes of Health.

## Conflict of interest

The authors declare that they have no conflicts of interest with the contents of this article.

## Abbreviations

A3A: APOBEC3A
A3B: APOBEC3B
A3BctdDM: APOBEC3B C-terminal domain double-mutant
A3BctdWT: APOBEC3B C-terminal domain wild-type
AMBER: APOBEC mediated base editing reporter
CBE: Cytosine base editor
dU: 2’-deoxyuridine
EndoQ: Pyrococcus furiosus endonuclease Q
FAM: 6-carboxyfluorescein
flA3BWT: Full-length APOBEC3B wild-type
FRET: Förster resonance energy transfer
IAB: Iowa Black® FQ
RADD: Real-time APOBEC mediated DNA deamination
RFU: Relative fluorescence unit
UDG: uracil-DNA glycosylase

## Notes

### Competing Interest Statement

The authors have declared no competing interest.

