## Supporting Information for "A real-time biochemical assay for quantitative analyses of APOBEC-catalyzed DNA deamination"

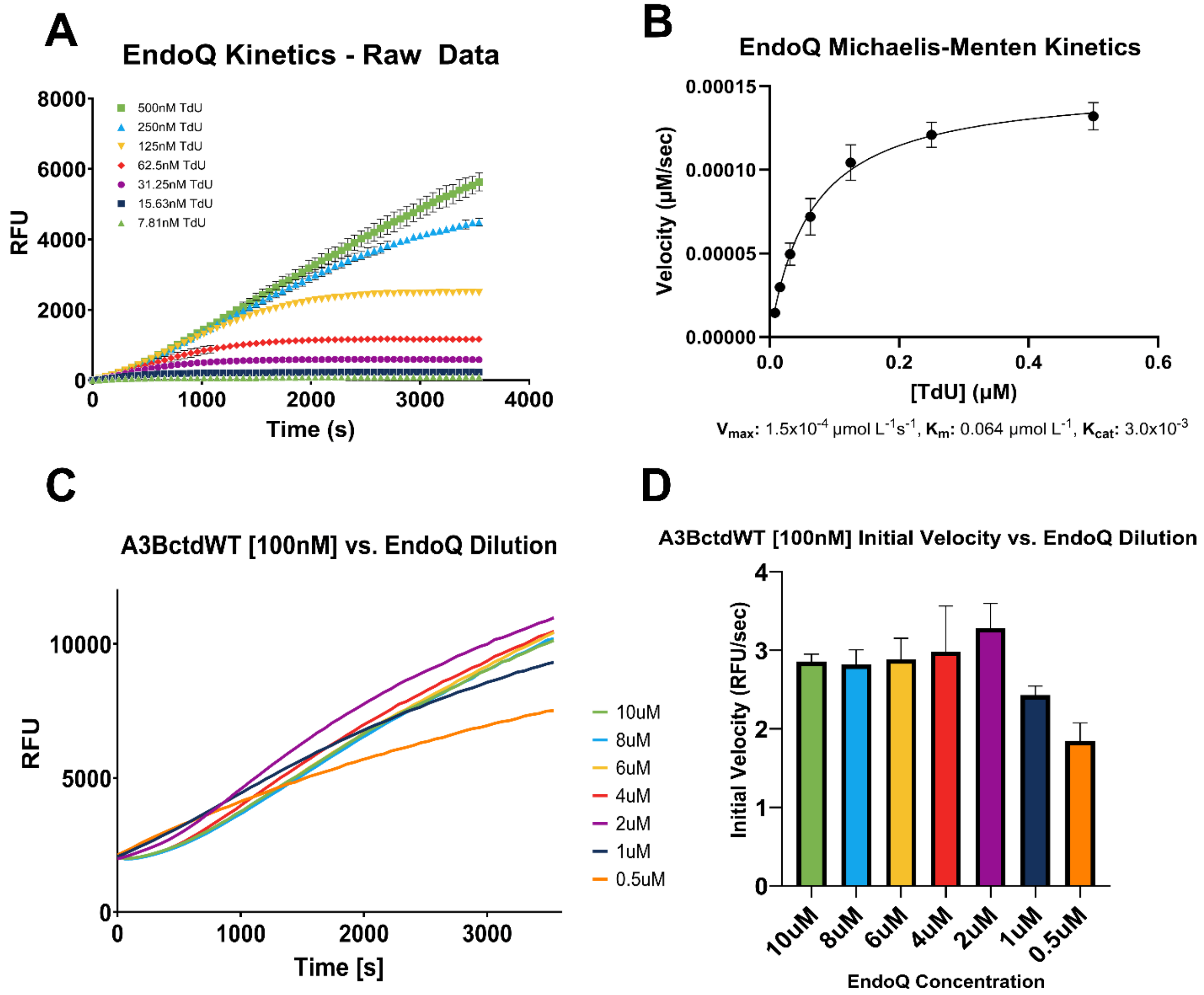


**Figure S1. Ensuring A3B was the rate-limited enzyme in the RADD reactions. (A)** To determine the kinetic parameters of EndoQ, 50 nM enzyme was evaluated against a ½ dilution series of FAM-GTdUA-IAB reporter starting at 500 nM (N = 3 for each condition). **(B)** The maximum slope (initial velocity) for each condition from **A** was determined and converted to µM / second before fitting with the *Michaelis-Menten* model ($R^2$ = 0.98). The $V_{max}$ was determined to be $1.5x10^{-4}$ µmol $L^{-1}$ $s^{-1}$ 95% CI [$1.4x10^{-4}$ to $1.6x10^{-4}$ µmol $L^{-1}$ $s^{-1}$]. The $K_M$ was determined to be 0.064 µmol $L^{-1}$ 95% CI [0.052 to 0.079 µmol $L^{-1}$] and the $k_{cat}$ of the enzyme was determined to be $3.0x10^{-3}$ $s^{-1}$ 95% CI [$2.8x10^{-3}$ to $3.2x10^{-3}$ $s^{-1}$]. (**C**, **D**) A dilution series of EndoQ was then evaluated in reactions containing the FAM-GTCA-IAB reporter and 100 nM A3BctdWT. The results showed that 2 µM EndoQ provided the optimal signal readout and that concentrations higher than 2 µM began to show decreased fluorescence signal. The kinetic traces in **(C)** depicts the averaged fluorescence readout of each EndoQ concentration condition while **(D)** depicts the slope (representing initial reaction velocity) of each trace.

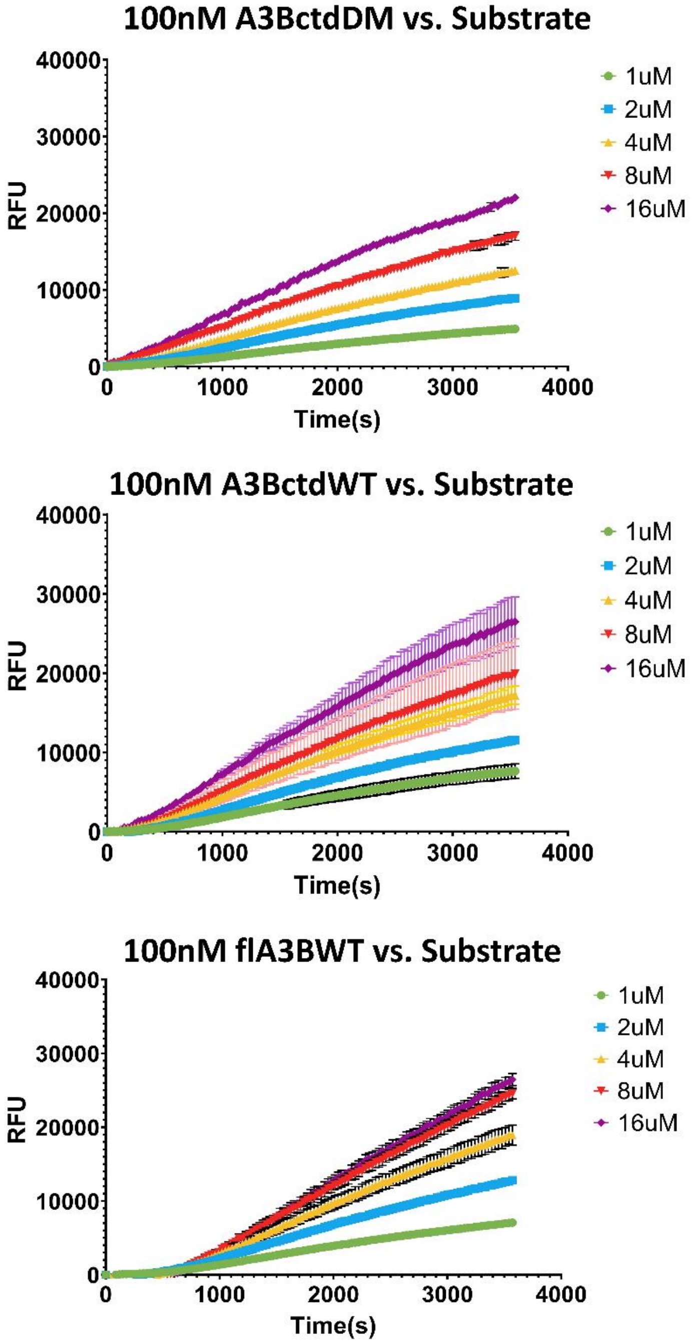


**Figure S2. Real-time fluorescence read-out of data used to calculate the enzyme kinetics shown in Fig. 3 (N = 3 for each reporter concentration).** Each reaction contained 100 nM of the respective APOBEC3B variant, 2 μM EndoQ and 1-16 μM FAM-GTCA-IAB substrate. Each reaction was prepared in parallel with a background control containing no APOBEC3B, 2 μM EndoQ, and the corresponding reporter concentration. These background controls were subtracted from the traces displayed above.

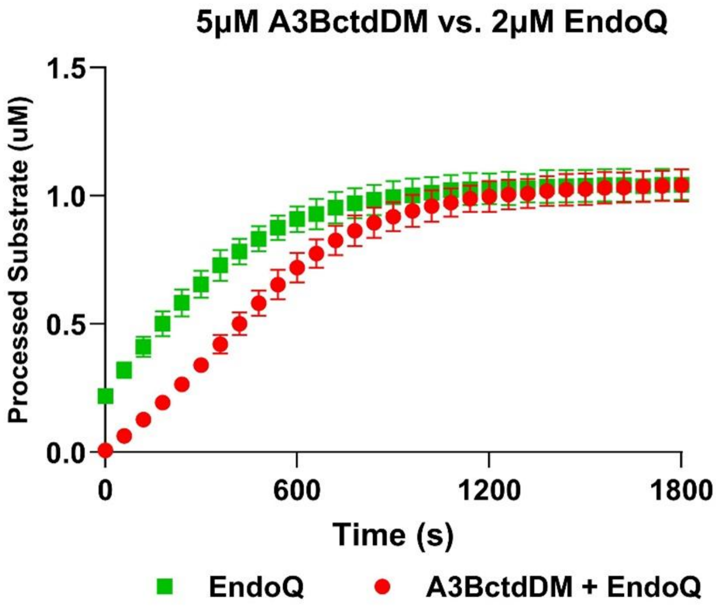


**Figure S3. Real-time fluorescence read-out of 5 μM A3BctdDM and 1 μM TC reporter.** The trace in (red) depicts the real-time readout of 5 μM A3BctdDM in the presence of 1 μM TC reporter oligo and 2 μM EndoQ, while the trace in (green) represents 2 μM EndoQ control in the presence of 1 μM TdU reporter (N = 3 for each condition). The EndoQ control data was used to calculate $k_{obs}$ / $k_{cat}$ in **Fig. 4.** The A3BctdDM data was used to fit the sequential enzyme reaction model in **Fig. 4** and subsequently estimate $k_{cat}$.

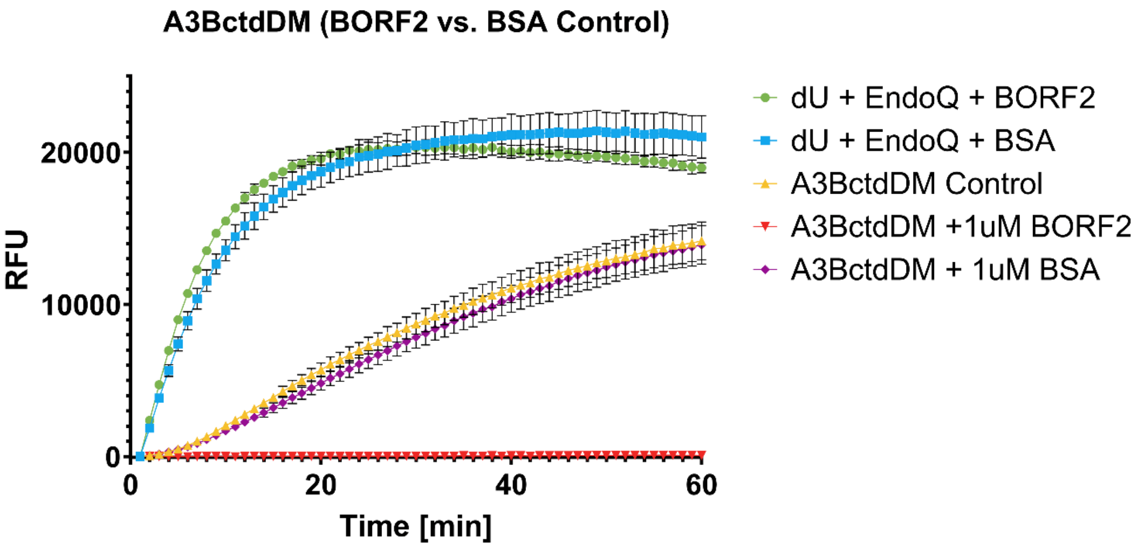


**Figure S4. Examining potential effects of protein crowding on the RADD assay.** Both 500 nM A3BctdDM and 2 μM EndoQ were tested against either 1 μM BORF2 and 1 μM BSA (N = 3). When combined with BORF2, A3BctdDM was completely inhibited, but when combined with an equal concentration of BSA, showed no significant difference in activity compared to the control. A similar trend was observed for EndoQ, with the exception of increased signal stability over time in the presence of BSA.

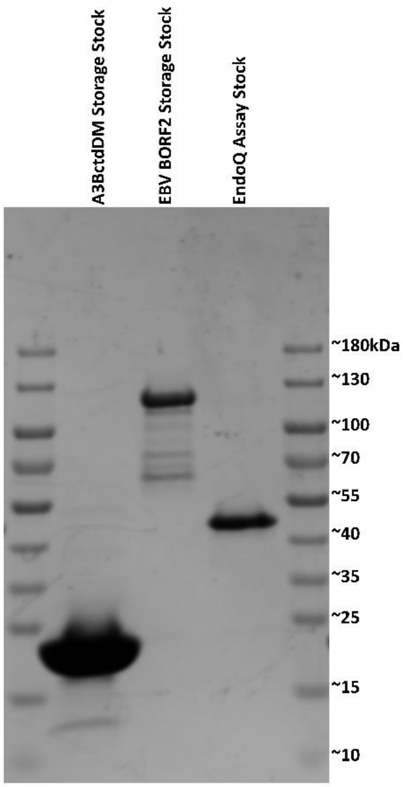


**Figure S5. SDSPage gel of enzymes used in the BORF2 inhibition RADD assay.** Both A3BctdDM and BORF2 were thawed from storage stocks and diluted before each experiment, while EndoQ was thawed from pre-diluted storage stocks. Both A3BctdDM and EndoQ are >95% pure, however BORF2 showed significant contamination. The purity of BORF2 was estimated via band quantification and found to be ~62% pure. This metric was used to adjust all future nanodrop measurements for this stock.

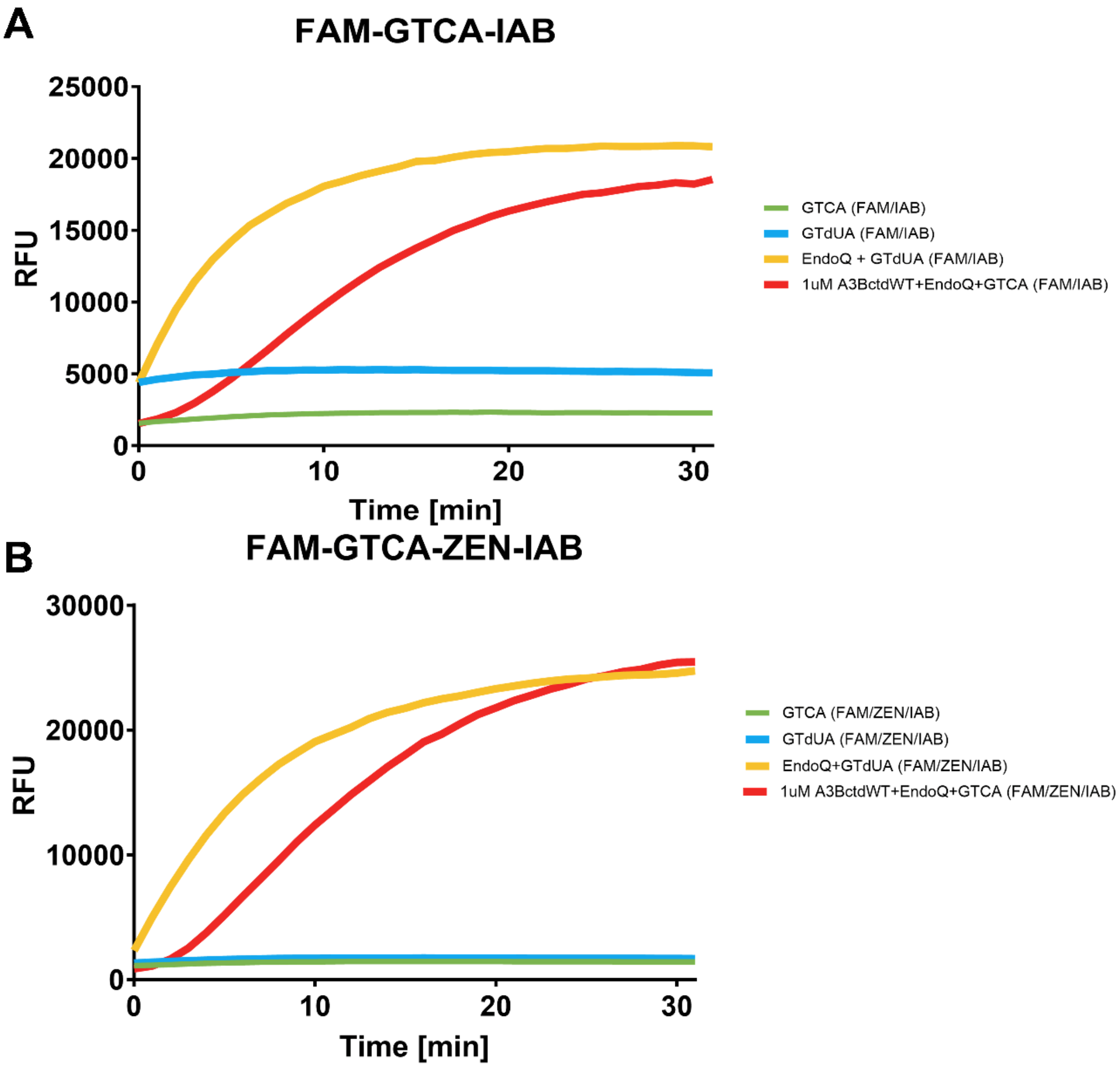


**Figure S6. Using internally quenched reporter oligos for improved signal and reduced background in the RADD assay. (A)** RADD assay using 1 μM standard reporter and control oligos (FAM-TC and TdU-IAB) versus 1 μM A3BctdWT. As previously observed, TdU shows a naturally high background signal which requires additional processing to normalize with the TC oligo background, there is also a small, but noticeable increase in the negative control signals between 0 and 5 minutes before plateauing. **(B)** RADD assay using 1 μM doubly quenched reporter and control oligos (FAM-TC and TdU-ZEN-IAB). This reporter system completely eliminates the high background seen with the TdU reporter, and largely eliminates the signal increase in the negative controls. This reporter system also produced both lower signal from negative controls and greater signal from positive controls.

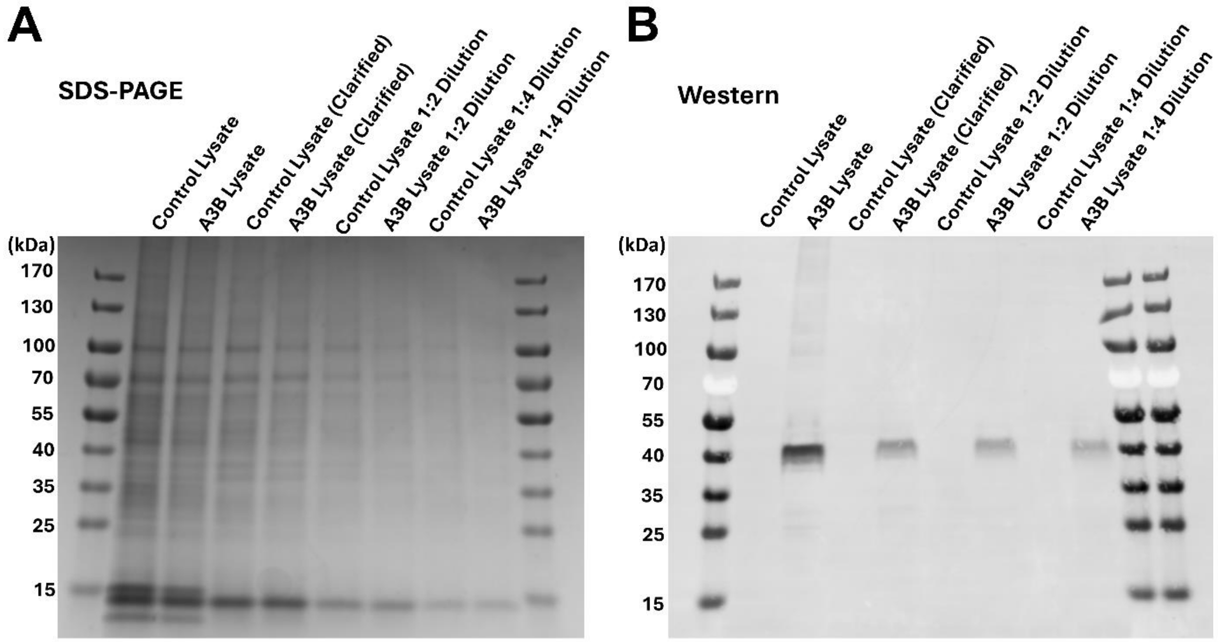


**Figure S7. SDS-PAGE and Western blot controls for cell lysate RADD. (A)** SDS-PAGE gel showing the total protein present in the raw lysate as well as the clarified lysate and diluted samples used in **Fig. 6** from both control and A3B-expressing cells. **(B)** Western blot of identical samples to those seen in the SDS-PAGE gel, blotted with α-HA rabbit primary and goat α-rabbit 680TL secondary antibodies. HA-A3B is clearly present in all A3B lysate lanes while no signal was detected in the control lanes. As expected, each dilution shows a gradual decrease in the HA-A3B signal.

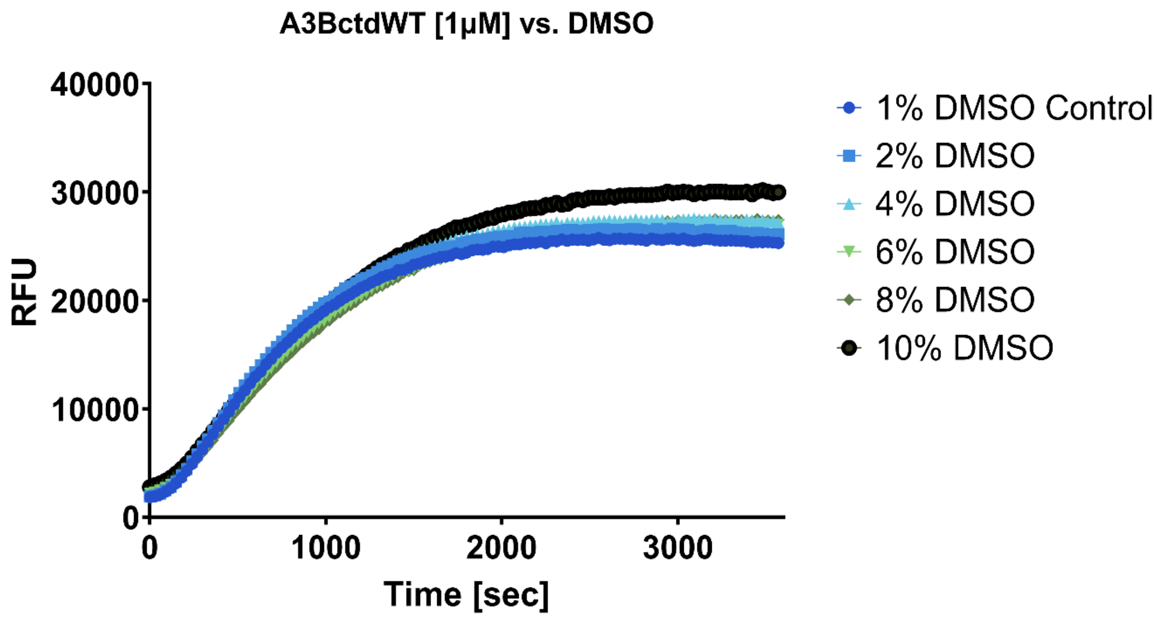


**Figure S8. Examining the effects of DMSO concentration on the RADD assay.** Using 1 μM A3BctdWT, there are not observable negative effects on the enzyme's ability to deaminate the reporter substrate. Interestingly, there appears to be a trend of increased fluorescence signal correlated with the increase in DMSO concentration. Small molecule inhibitors are often solubilized in DMSO, making it imperative that the assay is capable of functioning under relatively high DMSO concentrations. These findings indicate that the RADD assay is viable for use in small molecule inhibitor screens against A3B.
